# Yellow fever virus spread in Rio de Janeiro and Espírito Santo, 2016-2019: Phylodynamic assessment to improve intervention strategies

**DOI:** 10.1101/711994

**Authors:** Marta Giovanetti, Marcos Cesar Lima de Mendonça, Vagner Fonseca, Maria Angélica Mares-Guia, Allison Fabri, Joilson Xavier, Jaqueline Goes de Jesus, Tiago Gräf, Cintia Damasceno dos Santos Rodrigues, Carolina Cardoso dos Santos, Simone Alves Sampaio, Flavia Lowen Levy Chalhoub, Fernanda de Bruycker Nogueira, Julien Theze, Alessandro Pecego Martins Romano, Daniel Garkauskas Ramos, Andre Luiz de Abreu, Wanderson Kleber Oliveira, Rodrigo Fabiano do Carmo Said, Carlos F. Campelo de Alburque, Tulio de Oliveira, Carlos Augusto Fernandes, Shirlei Ferreira Aguiar, Alexandre Chieppe, Patrícia Carvalho Sequeira, Nuno Rodrigues Faria, Rivaldo Venâncio Cunha, Luiz Carlos Junior Alcantara, Ana Maria Bispo de Filippis

**Affiliations:** Laboratório de Flavivírus, Instituto Oswaldo Cruz Fiocruz, Rio de Janeiro, Brazil; Laboratório de Genética Celular e Molecular, ICB, Universidade Federal de Minas Gerais, Belo Horizonte, Minas Gerais, Brazil; KwaZulu-Natal Research Innovation and Sequencing Platform (KRISP), College of Health Sciences, University of KwaZuluNatal, Durban 4001, South Africa; Instituto Gonçalo Moniz, Fundação Oswaldo Cruz, Ministério da Saúde, Salvador-BA; Department of Zoology, University of Oxford, OX1 3PS, UK; Coordenação Geral de Vigilância de Arboviroses (CGARB); Coordenação Geral dos Laboratórios de Saúde Pública/Secretaria de Vigilância em Saúde, Ministério da Saúde, (CGLAB/SVS-MS) Brasília, Distrito Federal, Brazil; Secretaria de Vigilância em Saúde, Ministério da Saúde (SVS-MS), Brasília, Distrito Federal, Brazil; Organização Pan-Americana da Saúde/Organização Mundial da Saúde, Brasília, Distrito Federal, Brazil; Laboratório Central de Saúde Pública Noel Nutels (LACEN-RJ), Rio de Janeiro, Brazil; Superintendência de Vigilância do Estado, Rio de Janeiro, Brazil; Fundação Oswaldo Cruz, Bio-Manguinhos, Rio de Janeiro, RJ, Brasil; Universidade Federal do Mato Grosso do Sul, Faculdade de Medicina, Campo Grande, MS, Brazil

**Keywords:** Yellow fever, outbreak, southeast Brazil, genomic surveillance, outbreak response

## Abstract

The recent re-emergence of yellow fever virus (YFV) in Brazil has raised serious concerns due to the virus’ rapid dissemination in the southeastern region. To better understand YFV genetic diversity and dynamics during the recent outbreak in southeastern Brazil we generated 18 complete and near-complete genomes from the peak of the epidemic curve from non-human primates (NHPs) and human infected cases across Espírito Santo and Rio de Janeiro states. Genomic sequencing of 18 YFV genomes revealed the timing, source and likely routes of yellow fever virus transmission and dispersion during the one of the largest outbreaks ever registered in Brazil. We showed that the recent YFV epidemic spillover southwards several times from Minas Gerais to Espírito Santo and Rio de Janeiro states in 2016 to 2019. The quick production and analysis of data from portable sequencing could identify the corridor of spread of YFV. These findings reinforce that real-time and continued genomic surveillance strategies can assist in the monitoring and public health responses of arbovirus epidemics.

**IMPORTANCE:** Arbovirus infections in Brazil including Yellow Fever, Dengue, Zika and Chikungunya result in considerable morbidity and mortality and are pressing public health concerns. However, our understanding of these outbreaks is hampered by limited availability of real time genomic data. In this study, we investigated the genetic diversity and spatial distribution of YFV during the current outbreak in southeastern Brazil. To gain insights into the routes of YFV introduction and dispersion, we tracked the virus by sequencing YFV genomes sampled from non-human primates and infected patients from the southeastern region. Our study provides an understanding of how YFV initiates transmission in new Brazilian regions and illustrates that near-real time genomics in the field can augment traditional approaches to infectious disease surveillance and control.

## INTRODUCTION

Yellow fever (YF) is a vector-borne disease that is endemic in tropical areas of Africa and South America (1). The aetiologic agent is the yellow fever virus (YFV), a single-stranded positive sense, RNA virus belonging to the *Flaviviridae* family (2). YFV diversity can be classified into four distinct genotypes, which have been named based on their geographical distribution: East African, West African, South American I, and South American II genotypes (3–6).

In the Americas, YFV transmission can occur via two main epidemiological transmission cycles: the sylvatic (or jungle) and the urban (domestic) cycles. In the sylvatic cycle non-human primates (NHPs) are infected through the bite of mosquito vectors such as *Haemagogus spp.* and *Sabethes spp.* (7, 8). However, in the urban cycle, humans can be infected by *Aedes spp.* mosquitoes biting (9). YFV infection in humans shows a wide spectrum of disease severity including asymptomatic infection, mild illness with dengue-like symptoms, including fever, nausea, vomiting and fatigue, and severe disease, including fever with jaundice or hemorrhage and death (10).

While eradication is not feasible due to the wildlife reservoir system, large-scale vaccination coverage provides considerable protection against the re-urbanization of YFV transmission (11). However, despite the availability of effective vaccines, YF remains an important public health issue in Africa and South America. In late 2016, a severe re-emergence of YFV epidemic has been reported in southeastern Brazil. The epidemic has evolved to become the largest observed in the country in decades, reaching areas close to the Atlantic rainforest (11, 12). YFV 2016-2017 epidemic in Brazil accounted for 1,412 epizootics, 777 YF human confirmed cases, most of which in southeast Brazil (Minas Gerais n=465; Sao Paulo n=22, Rio de Janeiro n=25; Espírito Santo n=252 confirmed cases), and 261 human deaths (13). Following this epidemic new cases were reported between 2017-2018 and in that period 864 epizootics, 1,376 YF human confirmed cases and 483 human deaths were registered, with the southern states among the most affected by the YFV epidemic (Minas Gerais n=532; Sao Paulo n=377, Rio de Janeiro n=186; Espírito Santo n=6 confirmed cases) (14). The epidemic persisted in 2018-2019 and accounted for 1,883 NHP notified cases (n=20 confirmed NHP cases) and 12 human confirmed cases, including 5 human deaths from the state of São Paulo. Most of the confirmed epizootic cases was registered in the southeastern states (95%) (São Paulo (n=10); Rio de Janeiro (n=8) and Minas Gerais (n=1) (13–15).

Although there is currently no evidence that urban transmission has occurred, the outbreak affected areas highly infested by *Ae. aegypti* and *Ae. Albopictus* where yellow fever vaccination was recently introduced in routinely of immunization program. This condition or behavior raises concern that, for the first time in decades, there might be high risk of YFV urban transmission in Brazil (16). New surveillance and analytical approaches are therefore needed to monitor this threat in real time.

Even so, there is limited information from genomic surveillance studies about the genomic epidemiology and the dissemination dynamics of 2016-2019 YFV circulating in Southeast Brazil. Previous studies have shown the spatial and evolutionary dynamics of the current YFV outbreak in different southeastern states, (11) and shed light regarding the possible co-circulation of distinct YFV lineages (17). Nevertheless, there is still limited information about the genomic epidemiology of YFV circulating in Espírito Santo and Rio de Janeiro states from genomic surveillance studies, impairs our understanding of the virus re-introduction, establishment and dissemination in those regions.

Thus, to better understand the re-emergence of the recent YFV epidemic in those regions, we analyzed a larger and updated dataset of recently released data of the YFV 2016-2019 epidemic in Brazil, including 18 newly generated complete genomes from human and NHPs from the Southeast states of Espírito Santo and Rio de Janeiro.

## RESULTS

### Molecular diagnostics and genome sequencing from clinical samples

Liver, spleen, kidney and blood samples from 14 NHPs and liver and serum samples from 4 human infected cases collected in Rio de Janeiro and Espírito Santo states, Southeast Brazil, between January 2017 and April 2018, were tested for YFV RNA using the RT-qPCR assay (18, 19) at the Flavivirus Laboratory at FIOCRUZ Rio de Janeiro (LABFLA/FIOCRUZ).

Most confirmed cases in NHPs were from animals of the *Alouatta* genus (42.9%; 6 of 14), followed by *Callithrix* (35.7%; 5 of 14), *Sapajus* (7.1%; 1 of 14) and *Leontopithecus rosalia* (14.3%; 2 of 14). PCR cycle threshold (Ct) values were on average 12.23 (range: 7.2 to 22.4) (**Table 1**).

**Table 1.** Epidemiological data for the sequenced samples.

| ID | CT value | Sample Type | Host | Species | State | Municipality | Collection Date | Age | Sex |
| --- | --- | --- | --- | --- | --- | --- | --- | --- | --- |
| RJ182 | 8,2 | Liver | NHP | Alouatta Sp | RJ | São Sebastião Do Alto | 09/03/2017 | NA | M |
| RJ193 | 10,2 | Liver | NHP | Alouatta Sp | RJ | São Sebastião Do Alto | 27/03/2017 | 1 | M |
| RJ141 | 22,4 | Serum | Human | - | ES | Ibatiba | 24/01/2017 | 16 | M |
| RJ183 | 11,2 | Serum | Human | - | RJ | São Sebastião Do Alto | 12/03/2017 | 25 | M |
| RJ194 | 6,5 | Liver | NHP | Alouatta Sp | RJ | São Sebastião Do Alto | 27/03/2017 | 15 | F |
| RJ147 | 21,9 | Whole blood | NHP | Alouatta Sp | ES | Domingos Martins | 31/01/2017 | NA | NA |
| RJ173 | 15 | Whole blood | NHP | Cebus Sp | ES | Itarana | 09/02/2017 | NA | NA |
| RJ184 | 14,4 | Liver | Human | - | ES | Cariacica | 13/03/2017 | 65 | M |
| RJ213 | 8,1 | Liver | NHP | Callithrix Sp | RJ | Valença | 22/01/2018 | 5 | F |
| RJ186 | 10,9 | Liver | NHP | Alouatta Sp | ES | Guarapari | 06/03/2017 | NA | NA |
| RJ177 | 11,5 | Serum | Human | - | ES | Brejetuba | 16/02/2017 | 46 | M |
| RJ188 | 9,9 | Whole blood | NHP | Callithrix Sp | ES | Cariacica | 08/03/2017 | NA | NA |
| RJ201 | 13,4 | Liver | NHP | Callithrix Sp | RJ | Nova Iguaçu | 28/11/2017 | 2 | F |
| RJ219 | 11,2 | Kidney | NHP | Callithrix Sp | RJ | Angra Dos Reis | 05/02/2018 | NA | NA |
| RJ189 | 13,7 | Whole blood | NHP | Alouatta Sp | ES | Serra | 20/03/2017 | NA | F |
| RJ216 | 7,2 | Liver | NHP | Callithrix Sp | RJ | Duas Barras | 25/01/2018 | 10 | F |
| LABFLA09 | 22.1 | Spleen | NHP | Leontopithecus Rosalia | RJ | Silva Jardim | 24/04/2018 | NA | NA |
| LABFLA10 | 11.76 | Liver | NHP | Leontopithecus Rosalia | RJ | Silva Jardim | 24/04/2018 | NA | NA |

ID=study identifier; Ct=RT-qPCR quantification cycle threshold value; State= RJ-Rio de Janeiro; ES-
Espírito Santo; Municipality=Municipality of residence; F=Female; M=Male; NA= Not Available.

To investigate the source and transmission of YFV and the genetic diversity of the virus circulating in human and NHPs across Rio de Janeiro and Espírito Santo states, we used the MinION handheld nanopore sequencer to generate 18 complete and near complete genomic sequences (average coverage = 89.9%; **Table 2**) using a previously described MinION sequencing protocol (11, 20).

**Table 2.**
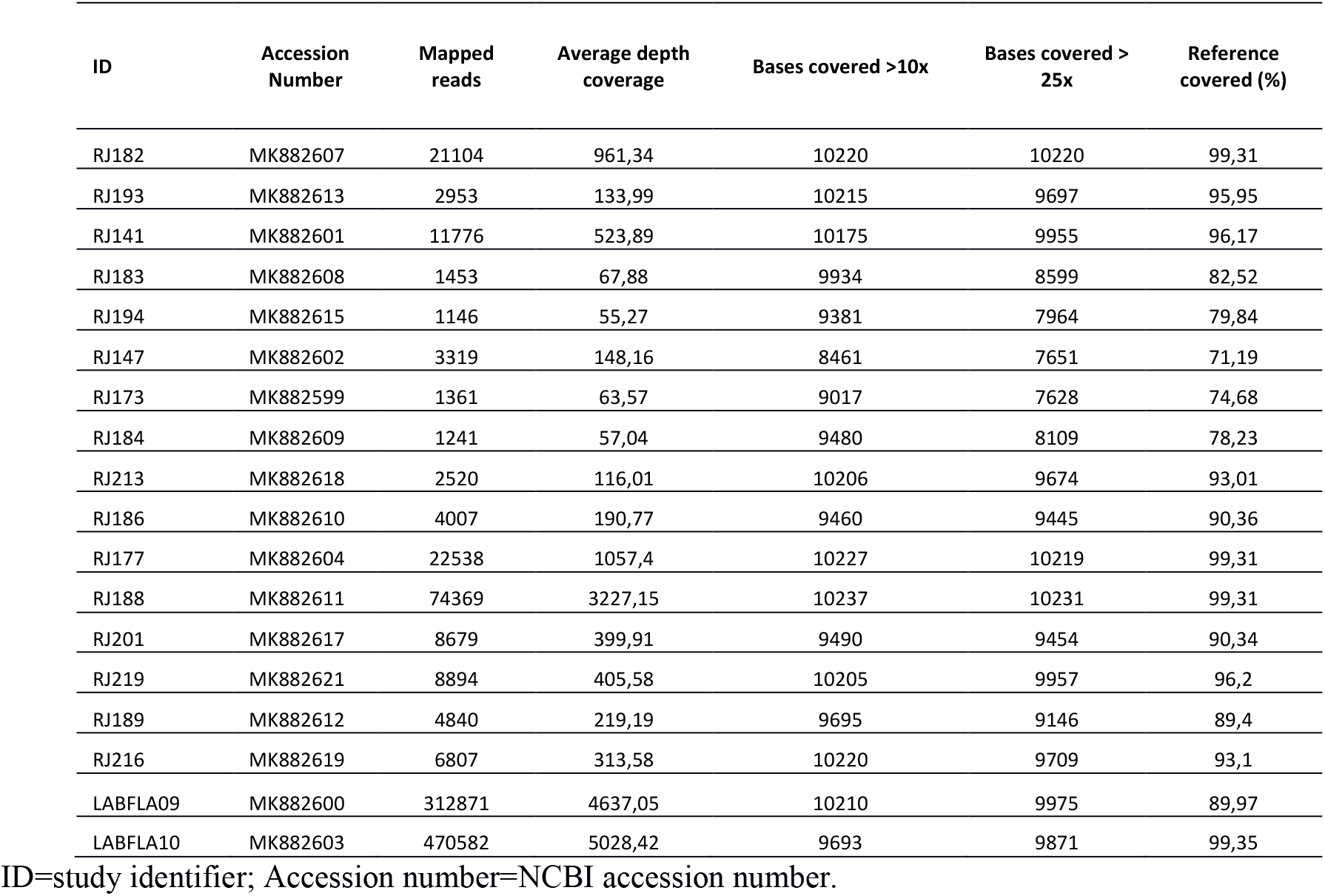
Sequencing statistics for the 18 new obtained sequences.

New sequences have been deposited in GenBank under accession numbers: MK882599- MK882604; MK882607-MK882613; MK882615; MK882617-MK882619; MK882621. YF samples sequenced in this study were geographically widespread across 6 municipalities of Rio de Janeiro and 7 municipalities of Espírito Santo, (**Figure 1A**).

**Figure 1.**
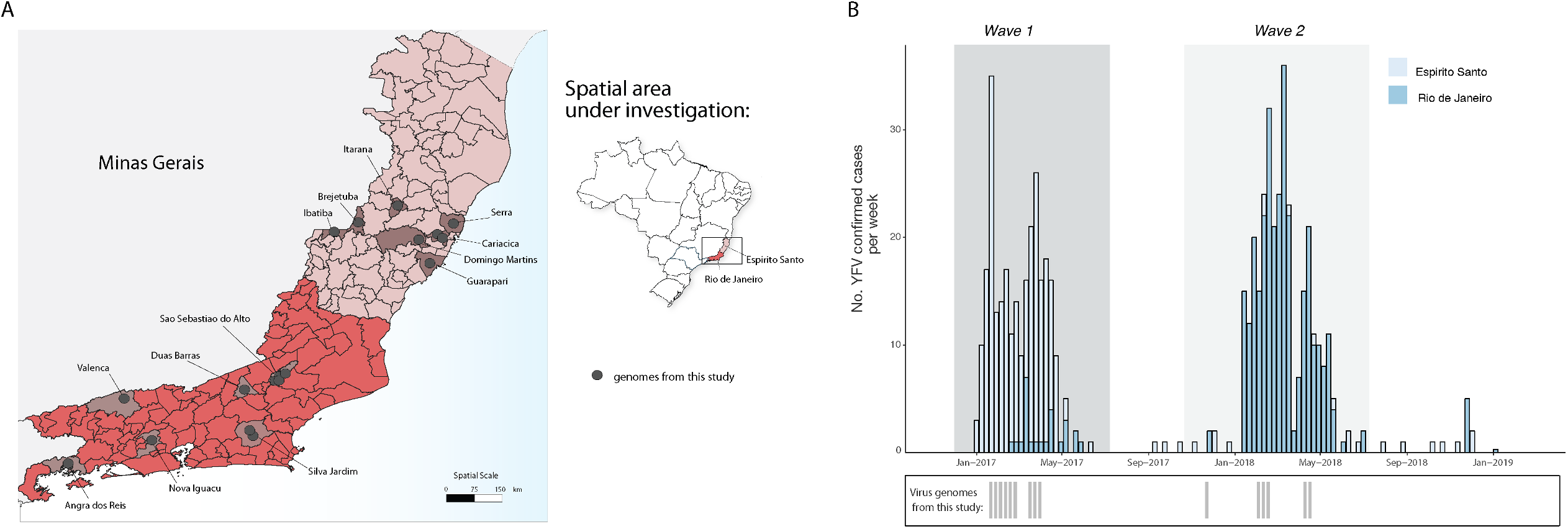
Spatial and temporal distribution of YF cases from Espírito Santo and Rio de Janeiro states during 2017 and 2019. **A**. Map of the states of Espírito Santo (ES) and Rio de Janeiro (RJ), located in south-eastern region of Brazil, and its municipalities. Circles indicate where samples from this study were collected. **B.** Time series of human (H) and Non-human primate YFV cases in ES and RJ states confirmed by serology, reverse transcription quantitative PCR (RT-qPCR), or virus isolation. Below, the dates of sample collection of the virus genomes generated in this study are shown in grey bars.

Figure 1 panel B shows the number of YFV confirmed cases in the Espirito Santo and Rio de Janeiro states respectively. Epidemiological data revealed two distinct YFV epidemic waves. The first epidemic wave (*wave 1*) is represented by the YFV cases mainly registered in the Espírito Santo state, during the first semester of 2017 (January to April, *n*= 252 cases), although some sporadic cases were reported in the following year (**Figure 1B**). The second wave (*wave 2*), in turn, is represented by YFV cases registered in Rio de Janeiro state during first semester of 2018 (February to May; *n*=220 cases) (**Figure 1B**). Although majority of cases in Rio de Janeiro occurred between February and March 2018, we can see that the re-emergence of YFV in that state was detected around March 2017, during epidemic wave 1 that mainly affected Espírito Santo state.

### Genetic history of YFV in Southeastern Brazil

To investigate the phylogenetic relationship of YFV strains circulating in the southeastern states of Espírito Santo and Rio de Janeiro we estimated a maximum likelihood (ML) phylogenetic tree for a dataset of 181 reference sequences comprising the four YFV lineages. Our ML phylogeny revealed that, as suspected, the newly generated YFV sequences belong to the South American I (SAI) lineage with high statistical support (bootstrap = 100%), clustering with other Brazilian isolates from the 2016-2019 epidemic (**Supplementary Figure 1**).

Subsequently, to investigate the dynamic of the YFV infection within the Southeast region, genetic analyses were conducted on a second dataset (dataset 2, *n* = 137), including recently published sequences from the YFV 2016-2019 epidemic in Brazil, belonging to the SA1 lineage. The time-scale of our phylogenetic estimates was consistent with recently studies (17, 21, 22) and confirmed the presence of two distinct lineages circulating in the current YFV epidemic, named hereafter as SA1 lineage 1 and SA1 lineage 2 (**Figure 2**). The SA1 lineage 1 comprises sequences from the northern and eastern regions of Minas Gerais, Bahia, Espírito Santo and Rio de Janeiro states and the time of the most recent common ancestor (TMRCA) of this lineage was dated back to September 2016 (95% BCI: July to November 2016). Meanwhile the SA1 lineage 2 comprises sequences from the southern municipalities of Minas Gerais state and sequences from the southeastern state of Sao Paulo and the TMRCA of this lineage was dated back around July 2016 (95% BCI: June to December 2016) (**Fig. 2**). Moreover, our time-scaled phylogeny showed that the sequences generated in this study clustered together with high support (pp=90 %) within SA1 lineage 1 (**Figure 2**).

**Figure 2.**
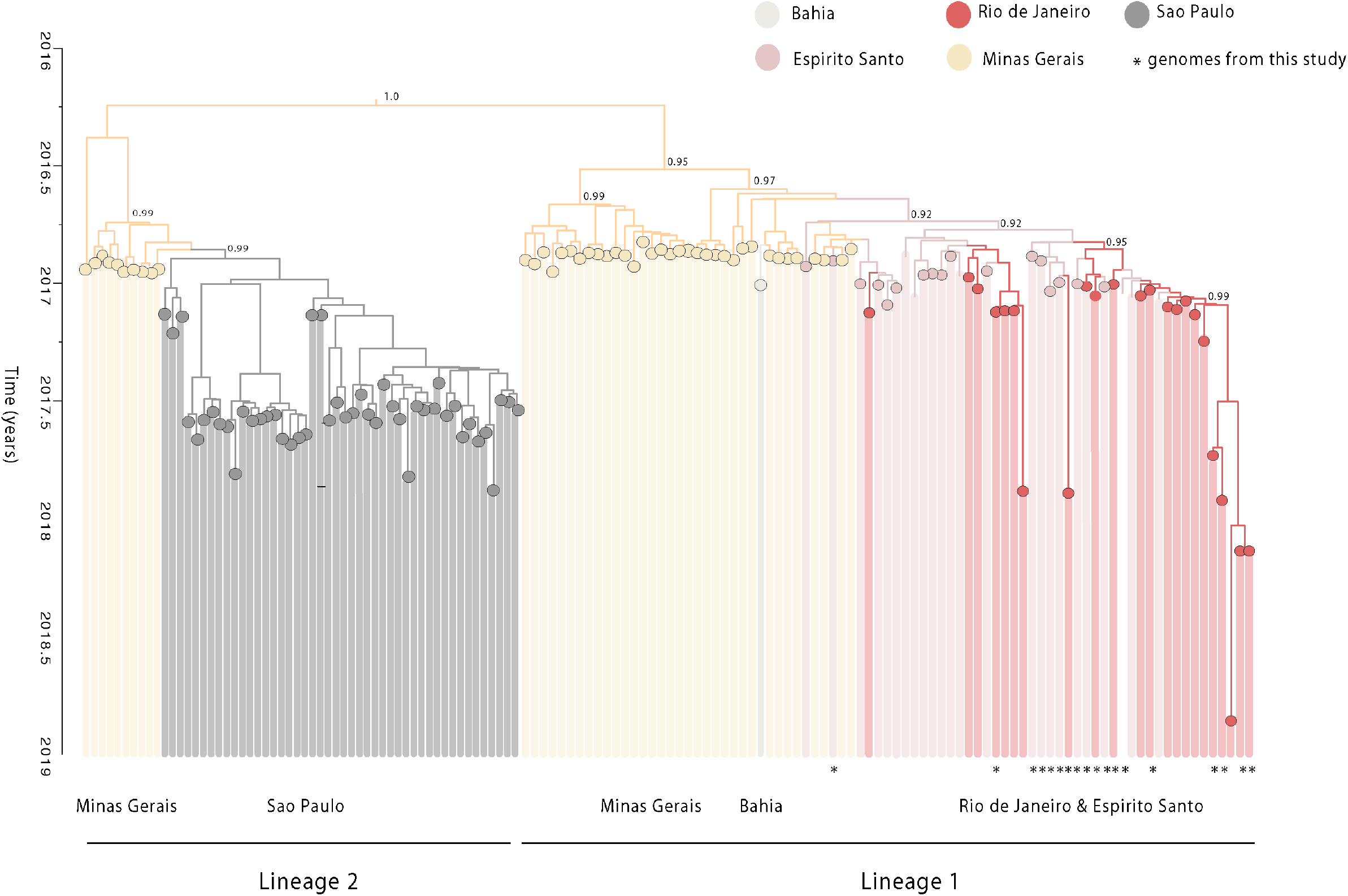
Time scaled phylogenetic tree of the current YF epidemic in Brazil. Molecular clock phylogeny obtained by combining the 18 new YFV complete genomic generated here (starred tips), plus public available data (n=137) of the YFV 2016-2019 epidemic in Brazil (11; 12; 17; 21-22). Numbers in nodes represent clade posterior probability >0.90. Branch colours represent different sampling locations.

In order to understand the transmission and the spatio-temporal evolution of the SA1 lineage 1, subsequently we analysed a subset of 81 (Dataset 3) sequences from this lineage (**Supplementary figure 2**). We performed a regression of genetic divergence from root to tip against sampling dates that confirmed sufficient temporal signal (r^2^=0.70) in this dataset. A time-scaled phylogenetic analysis using a Bayesian Markov Chain Monte Carlo (MCMC) framework (23) was then performed to investigate the time of introduction of the YFV into the Espírito Santo and Rio de Janeiro states (**Figure 3 A**). **Figure 3A** showed a zoom of our Bayesian time-scaled phylogeny highlighting the SA1 lineage 1 comprising the 2017-2019 YFV strains from Minas Gerais, Bahia, Espírito Santo and Rio de Janeiro states. Our analysis showed that samples from Espírito Santo were intermixed with sequences from Rio de Janeiro. This suggests that the YFV epidemic in Espírito Santo and Rio de Janeiro was not caused by a single introduction event, as observed in Sao Paulo (17, 21), but resulted from multiples introduction over time.

**Figure 3.**
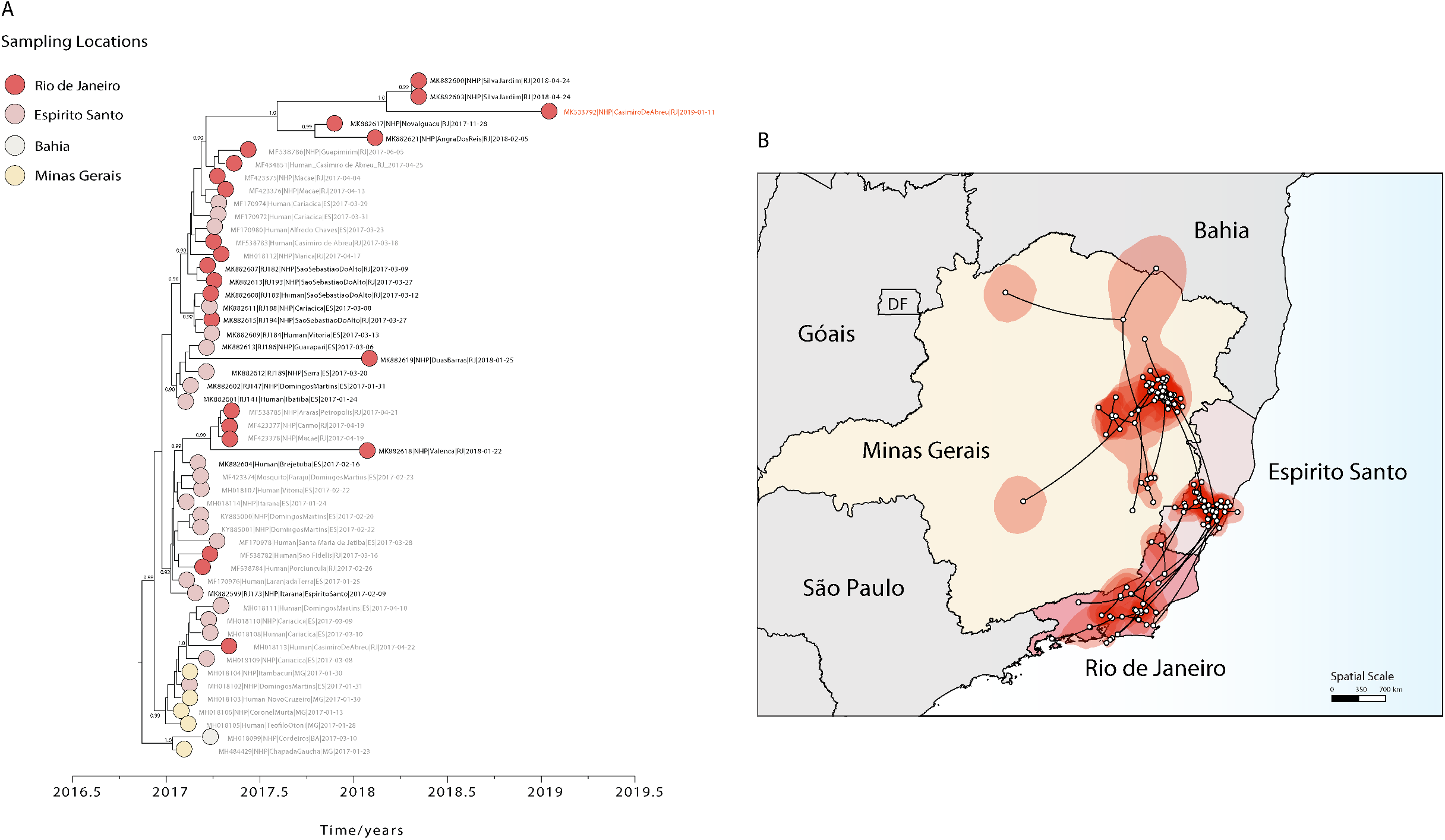
Spatio-temporal dynamics of the YFV SA1 lineage 1. **A**. Molecular clock phylogeny including the clade comprising the 2017-2019 YFV strains from Minas Gerais, Bahia, Espírito Santo and Rio de Janeiro states belonging to the SA1 lineage 1. Numbers along branches represent clade posterior probability >0.90. YFV isolates from Casimiro de Abreu, sampled in January 2019 is highlighted in red. Colours represent different locations. **B**. Reconstructed spatiotemporal continuous diffusion of the YFV SA1 lineage 1 outbreak clade. Phylogenetic branches are mapped in space according to the location of phylogenetic nodes (circles). Lines show the cross-state movement of the virus from Minas Gerais followed by movement to the states of Espírito Santo and Rio de Janeiro. Shaded regions show 95% credible regions of internal nodes.

We next used a continuous diffusion model to investigate how the SA1 lineage 1 has been spreading over space and time. We found evidence that YFV disseminated through southeastern Brazilian states using two distinct paths with an average dispersal rate of 0.12 km/day (95% HPD: 0.09 – 0.14 km/day). From the northern region of Minas Gerais state, YFV spread to the south region of Bahia state around January 2017 (95% BCI: December 2016 to February 2017) (**Figure 3**), and from the eastern region of Minas Gerais state YFV moved towards Espírito Santo state, (pp=0.99) with introductions estimated around January 2017 (95% BCI: November 2016 to January 2017) (**Figure 3 Panels A, B**). Since its introduction in the Espírito Santo state, the virus widespread through the neighboring state (**Figure 3**). Our analyses revealed that YFV was likely introduced in Rio de Janeiro state several times between January (95% BCI: December 2016 to February 2017) and March 2017 (February 2017 to May 2017), spreading southward from the border with Espírito Santo state and reaching Angra dos Reis municipality, which is located in the southern region of Rio de Janeiro. Our data further suggests that after its first introduction in Rio de Janeiro the virus persisted until 2019, as indicated by the isolate MK533792 sampled, in the municipality of Casimiro de Abreu, in January 2019 (12) (**Figure 3 Panel A**).

## DISCUSSION

In this study, we generated and analysed 18 new YFV complete and near-complete genomic sequences from samples from humans and non-human primates collected in several municipalities in Espírito Santo and Rio de Janeiro states, in 2017-2018.

Despite previous studies have already shown the spatial and evolutionary dynamics of the current YFV outbreak in Brazil (11, 12, 17, 21), the shortage of genomic data from Espirito Santo and Rio de Janeiro states hampered the possibility of shed light on the re-emergence, establishment and the corridor of spread of the YFV transmission in those regions.

The generated genomic data provides a more detailed understanding of the introduction and progression of YFV SA1 lineage 1 and reveals the timing, source and likely routes of yellow fever virus transmission and dispersion during the largest outbreak in Brazil in decades.

According to the Ministry of Health epidemiological bulletin, the YFV re-emergence in the states of Espírito Santo and Rio de Janeiro was confirmed in these states in January and February 2017 respectively (13–15).

Our estimates indicated that YFV strains from the epidemic first emerged in Espírito Santo state from Minas Gerais around January 2017 (95% BCI: November 2016 to January 2017), which is consistent with epidemiological data (13–15). From the state of Espírito Santo, YFV spread southwards to the great metropolitan area of Rio de Janeiro state. Moreover, our data indicated that the circulation of YFV in Rio de Janeiro may have resulted from multiple and independent introductions events from Espírito Santo state, highlighting a complex dispersion dynamic of the current YFV outbreak in Brazil occurred between January (95% BCI: December 2016 to February 2017) and March 2017 (February 2017 to May 2017). Our data further suggest that after its first introduction in Rio de Janeiro the virus persisted until 2019, as indicated by the isolate MK533792 sampled in the municipality of Casimiro de Abreu in January 2019 (12). This estimation suggests the YFV have persisted in Rio de Janeiro state for approximately 24 months. This suggest that Rio de Janeiro state possesses the ecological conditions to maintain YFV outside the period of transmission (Dec to May) (12). Ultimately, given the abundance of sylvatic competent vectors (12) and non-human primates (21, 22), this data could indicate that there is some potential for the establishment of an enzootic transmission cycle of yellow fever in Mata Atlantica.

Epidemiological data also indicated two distinct YFV epidemic waves (13, 14). The first epidemic wave is represented by the YFV cases mainly registered in Minas Gerais and Espírito Santo state during the first semester of 2017, while the second wave is represented by the YFV cases registered in Rio de Janeiro state during first semester of 2018. Transmission of YFV in areas with susceptible NHPs species typically occurs in time periods characterized by environmental conditions suitable to support higher mosquito abundance (12, 24).

As previously suggested (17), we found evidence regarding the circulation of two distinct YFV lineage, that might have been spread to distinct evolutionary and diffusion rates. Using YFV genetic data, we estimate that the YFV SA1 lineage 1 spread at rate of 0.12 km/day (95% HPD: 0.09 – 0.14 km/day), that is slightly lower than previously estimates (11, 17). The substantial difference between previously estimates (11, 17) and ours might reflect the larger dataset analyzed in this study, that might explain differences in the rate of YFV spread among different areas as well as different lineage.

These findings reinforce that continued genomic surveillance strategies are needed to assist in the monitoring and understanding of arbovirus epidemics, which might help to attenuate public health impact of infectious diseases.

In this study we also demonstrate that by analyzing heterochronous datasets with samples collected in different time points and/or locations, phylodynamics becomes a powerful tool to prevent and identify the viral lineage movement, describe trends in epidemic spread and to improve intervention strategies (11, 25, 26).

Continued surveillance in human and non-human primates (NHP) in non-epidemic periods in the southeast region will be important in order to quantify the risk of new outbreaks and the establishment of new YFV transmission cycles in the region. In conclusion, our study shows that genomic data generated by real time portable sequencing technology can be employed to assist public health services in monitoring and understanding the diversity of circulating mosquito-borne viruses.

## MATERIALS AND METHODS

### Sample collection

Human and non-human primate samples were collected, under the guidelines of a national strategy of YF surveillance, for molecular diagnostics by the Flavivirus Laboratory (LABFLA) at Oswaldo Cruz Foundation (Fiocruz) in Rio de Janeiro, Brazil, which is a Brazilian Ministry of Health Regional Reference Laboratory for arboviruses. The majority of samples were linked to a digital record that collated epidemiological and clinical data such as date of sample collection, municipality of residence, neighborhood of residence, demographic characteristics (age and sex) and date of onset of clinical symptoms.

### Ethical statement

The project was supported by the Pan American World Health Organization (PAHO) and the Brazilian Ministry of Health (MoH) as part of the arboviral genomic surveillance efforts within the terms of Resolution 510/2016 of CONEP (Comissão Nacional de Ética em Pesquisa, Ministério da Saúde; National Ethical Committee for Research, Ministry of Health). The diagnostic of YFV infection at LABFLA was approved by the Ethics Committee of the Oswaldo Cruz Institute CAAE90249218.6.1001.54248.

### RT-qPCR

Total RNA was extracted from tissue and serum samples using MagMAX™ Pathogen RNA/DNA kit (Life Technologies^TM^, Carlsbad CA, USA) in accordance with the manufacturer’s instructions. Viral RNA was detected using two previously published RT-qPCR techniques (18, 19).

### cDNA synthesis and whole genome nanopore sequencing

Sequencing was attempted on the 18 selected RT-PCR positive samples regardless of Ct value as previously described (11, 20, 26). All positive samples were submitted to a cDNA synthesis protocol (11, 20) using ProtoScript II First Strand cDNA Synthesis Kit. Then, a multiplex tiling PCR was attempted using the previously published YFV primer scheme and 30 cycles of PCR using Q5 High-Fidelity DNA polymerase (NEB) as previously described (20). Amplicons were purified using 1x AMPure XP Beads (Beckman Coulter) and cleaned-up PCR products concentrations were measured using Qubit™ dsDNA HS Assay Kit on a Qubit 3.0 fluorimeter (ThermoFisher). DNA library preparation was performed using the Ligation Sequencing Kit (Oxford Nanopore Technologies) and the Native Barcoding Kit (NBD103, Oxford Nanopore Technologies, Oxford, UK). Sequencing library was generated from the barcoded products using the Genomic DNA Sequencing Kit SQK-MAP007/SQK-LSK208 (Oxford Nanopore Technologies). Sequencing library was loaded onto a R9.4 flow cell (Oxford Nanopore Technologies).

### Generation of consensus sequences

Consensus sequences for each barcoded sample were generated following a previously published approach (20). Briefly, raw files were basecalled using Albacore, demultiplexed and trimmed using Porechop, and then mapped with *bwa* to a reference genome (GenBank accession number JF912190). Nanopolish variant calling was applied to the assembly to detect single nucleotide variants to the reference genome. Consensus sequences were generated; non-overlapped primer binding sites, and sites for which coverage was <20X were replaced with ambiguity code N. Sequencing statistics can be found in **Table 1**. Accession numbers of newly generated sequences are: MK882599-MK882604; MK882607-MK882613; MK882615; MK882617-MK882619; MK882621.

### Collation of YFV complete genome datasets

Genotyping was first conducted using the phylogenetic Yellow fever typing tool available at http://www.krisp.org.za/tools.php. The genome sequences generated here were combined with a dataset comprising previously published genomes from the 2016-2019 YFV epidemic in Brazil (11,12, 17, 21, 22). Two complete or near-complete YFV genome datasets were generated. Dataset 1 (*n* = 199) comprised the data reported in this study (*n* = 18) plus (*n* = 181) complete or almost complete YFV genomic sequences (>10,000 bp), retrieved from NCBI in June 2019 and covering all four existing genotypes. Subsequently, to investigate the dynamic of the YFV infection within the Southeast region, genetic analyses were conducted on a smaller dataset (dataset 2) including a larger and updated dataset of recently released data of the YFV 2016-2019 epidemic in Brazil belonging to the SA1 lineage (*n* = 137). Thus, to understand the transmission and the spatio-temporal evolution of the YFV SA1 lineage 1, from this dataset, we generated a subset (dataset 3) that included all identified sequences from that lineage (*n* = 81). Maximum likelihood (ML) phylogenetic trees were estimated using RAxML (27) under a GTR + Γ_4_ nucleotide substitution model. Statistical support for phylogenetic nodes was estimated using a ML bootstrap approach with 1000 replicates.

In order to investigate the temporal signal in our YFV datasets 2 and 3 we regressed root-to-tip genetic distances from this ML tree against sample collection dates using TempEst v 1.5.1 (http://tree.bio.ed.ac.uk) (28).

### Dated phylogenetics

To estimate time-calibrated phylogenies dated from time-stamped genome data, we conducted phylogenetic analysis using a Bayesian software package (23). Here we used the GTR + Γ_4_ nucleotide substitution model and Bayesian Skygrid tree prior (29) with an uncorrelated relaxed clock with a lognormal distribution (30). Analyses were run in duplicate in BEASTv.1.10.4 (23) for 50 million MCMC steps, sampling parameters and trees every 5000^th^ step. A non-informative continuous time Markov chain reference prior on the molecular clock rate was used (31). Convergence of MCMC chains was checked using Tracer v.1.7.1 (32). Maximum clade trees were summarized using TreeAnnotator after discarding 10% as burn-in.

### Phylogeographic analyses

To investigate the spread of YFV in Southeast Brazil, we analysed in more detail the SA1 Lineage 1 that includes the n=81 sequences (**Figure 2**). We used a skygrid coalescent tree prior (33) and a continuous phylogeographic model that uses a relaxed random walk to model the spatial diffusion of lineages. Dispersal velocity variation among lineages was modelled using a Cauchy distribution (34, 35). Virus diffusion through time and space was summarized using 1000 phylogenies sampled at regular intervals from the posterior distribution (after exclusion of burn-in). Sampling location of each geo-referenced YFV sequences from Espírito Santo and Rio de Janeiro state are listed in **Supplementary table 1**. Georeferenced and time-stamped sequences were analysed in BEAST v.1.10.4 (23) using the BEAGLE library (36) to enhance computational speed.

## Data availability

New sequences have been deposited in GenBank under accession numbers: MK882599- MK882604; MK882607-MK882613; MK882615; MK882617-MK882619; MK882621.

## ACKNOWLEDGMENTS

The authors thank all personnel from Health Surveillance System from Rio de Janeiro that coordinated surveillance and helped with data collection and assembly. The authors also thank Ronaldo Lapa e Solange Regina Conceição from the Instituto Oswaldo Cruz (Fiocruz – RJ) for their support. This work was supported by the ZiBRA2 project supported by the Brazilian Ministry of Health (SVS-MS) and the Pan American Organization (OPAS) and founded by Decit/SCTIE/MoH and CNPq (440685/2016-8 and 440856/2016-7); by CAPES (88887.130716/2016-00, 88881.130825/2016-00 and 88887.130823/2016-00); by EU’s Horizon 2020 Programme through ZIKAlliance (PRES-005-FEX-17-4-2-33). The Flavivirus Laboratory activities was also supported by Faperj (Fundação Carlos Chagas Filho de Amparo à Pesquisa do Estado do Rio de Janeiro) under the grant no. E-26/2002.930/2016, by the International Development Research Centre (IDRC) Canada over the grant 108411-001 and Horizon 2020 through ZikaPlan and ZikAction under the grant’s agreement No. 734584 and 734857. MG is supported by Fundação de Amparo à Pesquisa do Estado do Rio de Janeiro - FAPERJ.

## Author’s Contributions

Conception and design: M.G., M.C.L.M., V.F., L.C.J.A.; and A.M.B.F. Investigations: M.G., M.C.L.M., V.F., J.G.J., J.X., M.A.M.G., A.F., C.D.S.R., C.C.S., S.A.S., F.L.L.C., F.B.N. Data Curation: M.G., V.F., T.G., J.T., N.R.F. L.C.J.A and A.M.B.F. Formal Analysis: M.G., M.C.L.M., and V.F. Writing – Original Draft Preparation: M.G., M.C.L.M., V.F., L.C.J.A and A.M.B.F. Revision: M.G., T.G., A.C., P.S.C., A.P.M.R., D.G.R., T.O., N.R.F., L.C.J.A and A.M.B.F. Resources: L.C.J.A, A.P.M.R., D.G.R., A.L.A., W.K.O., C.C.A., C.A.F., S.F.A., A.C., R.V.C., and A.M.B.F.

## Supplementary Information

**Supplementary Figure 1**. Molecular phylogenetics of the Brazilian YFV epidemic. Maximum likelihood phylogeny of complete YFV genomes showing the outbreak clade (gray triangle) within the South American I (SA1) genotype. The scale bar is in units of substitutions per site (s/s).

**Supplementary Figure 2**. Molecular clock phylogeny including the clade comprising the new isolates plus all the YFV strains from the 2017-2019 outbreak belonging to the SA1 lineage 1 clade. Numbers along branches represent clade posterior probability >0.90. Colours represent different locations.

